# DOME Copilot: A resource to automate transparent reporting of artificial intelligence methods

**DOI:** 10.64898/2026.04.16.718888

**Authors:** Gavin Farrell, Omar Abdelghani Attafi, Styliani-Christina Fragkouli, Ignacio Heredia, Saul Fernández Tobías, Melissa Harrison, Henning Hermjakob, Matt Jeffryes, Mahta Mehdiabadi, Marta Obregón Ruiz, Matt Pearce, Nikos Pechlivanis, Federica Quaglia, Alvaro López García, Fotis Psomopoulos, Silvio C.E. Tosatto

## Abstract

Artificial intelligence (AI) methods are transforming life science research and witnessing unprecedented adoption across literature. While AI is driving impactful discoveries, this rapid growth in application has inundated researchers with poorly described models and datasets which lack transparency, impeding reusability and eroding trust. The DOME Recommendations aimed to address this issue and established structured reporting guidelines to standardize AI methodology descriptions. However, the need to manually create these comprehensive disclosures imposed a major bottleneck, requiring substantial effort from authors to comply. To bridge this gap, we present DOME Copilot, an open-source, easily deployable system that automates the generation of transparent AI methodology reports supplementing publications. Evaluated against a benchmark of human-created annotations, DOME Copilot was determined to match or exceed manual annotation quality across the majority of reporting fields while reducing the creation time from hours to minutes. Ultimately, the system provides a novel and scalable solution to restore confidence in complex life science AI method publications.

## Introduction

In recent years, there has been a paradigm shift in the rate of adoption and integration of artificial intelligence (AI) in biological research. This change has stemmed from several distinct but synergistic alignments across various enabling factors. This includes the advancements in graphics processing units (GPUs) and their wider accessibility to researchers such as through government investments in AI factories (Lopez *et al*. 2025). In tandem, the success and penetration of the FAIR data (Wilkinson *et al*. 2016) and Open Science movements (UNESCO 2021) have boosted the quality and availability of biological data in public repositories. This union of computational hardware and FAIR data has in turn facilitated high-profile application success cases drawing wider interest in adoption of AI, such as AlphaFold (Jumper *et al*. 2021) which received Nobel Prize recognition (Callaway 2024). Finally, the accessibility of high-quality open training materials, online resources and coding tools such as frontier large language models (LLMs) (Nvidia 2026), has substantially decreased the expertise overheads needed to practically implement AI approaches as standard analysis methods. These factors have jointly lowered the barriers to adopting highly complex AI methodology in life sciences, and this growth in application can be visualized directly through the increase in mentions of ‘artificial intelligence’ and ‘machine learning’ (ML) in life science manuscripts across the Europe PubMedCentral (PMC) (Rosonovski *et al*. 2024) literature database (Fig. 1).

**Figure 1.**
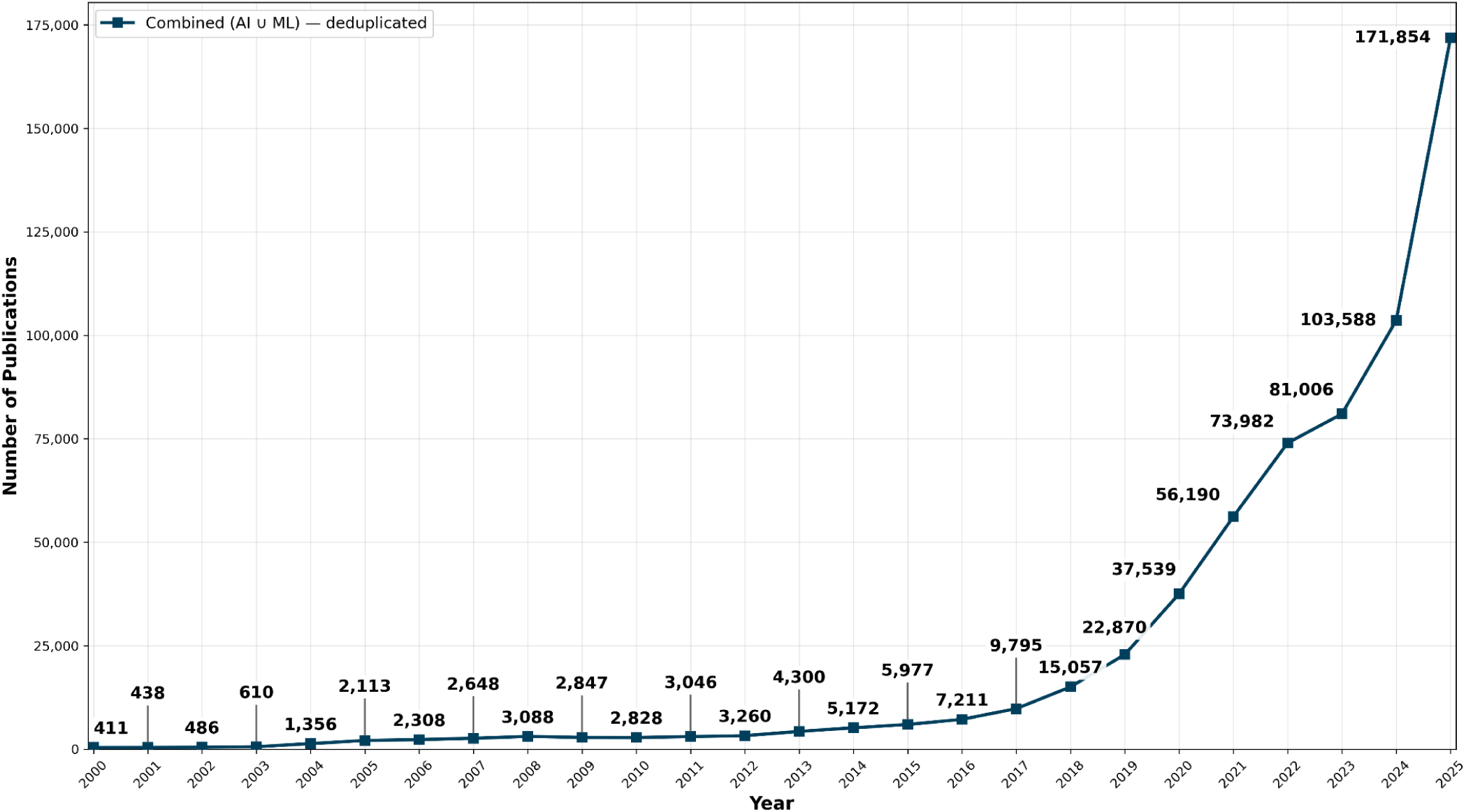
Europe PubMedCentral temporal growth of publications mentioning AI and ML. Data span from the year 2000 to the most recent full year (2025). The data points represent the number of publications containing the terms ‘artificial intelligence’ or ‘machine learning’ after deduplication where both are present.

Unfortunately, in spite of this rapid adoption rate and increase in breakthrough impactful methods ranging across diverse areas from drug discovery (Zhang K *et al*. 2025) to structural biology prediction (Abriata 2026), there has been a corresponding downside. These AI methods are often published without consideration for comprehensive or structured reporting, leading to obfuscated information impacting researchers’ ability to reuse, reproduce, and verify published results. Biologically focused journals, where AI approaches were previously not prevalent, are now seeing the proliferation of these methods, and domain-expert reviewers may be ill-equipped to evaluate key reporting nuances. This limitation stems from the technical complexity of what constitutes poor or improper model implementation. Consequently, a critical need has arisen for a solution similar to the FAIR data movement but tailored for AI methodology both within and beyond the life sciences. This growing crisis has been exemplified by landmark papers highlighting how gaps in structured reporting undermine literature trust, reusability, and reproducibility (Carter *et al*. 2019, Beam, Manrai, and Ghassemi 2020).

In response, several efforts have yielded ‘FAIR for AI’ guidelines or checklists, varying widely across depth, reporting criteria, structure, and targeted areas of domain application (Farrell *et al*. 2026). Gaining significant real-world adoption since its release, the DOME Recommendations (Walsh *et al*. 2021) have emerged as a flagship framework for structured reporting, driven by community uptake and integration into journal publishing workflows. DOME is a checklist of 21 key reporting fields across the four major methodological areas that constitute its acronym: Dataset, Optimization, Model, and Evaluation. It is directly scoped to implement standardized reporting for AI methods in publications to boost transparent information sharing. The framework has seen successful integration and attention such as within specific scientific communities (Palmblad *et al*. 2022), and in more formal settings like publisher workflows (Edmunds *et al*. 2026). However, despite seeing integration support by journals, a major obstacle to widespread adoption remained: the requirement for a dedicated infrastructure to easily create, host, and disseminate these completed DOME reports as a supplementary content type in a FAIR manner.

To resolve this infrastructure gap for these structured AI methodology reports, ELIXIR Europe (Harrow *et al*. 2021) funded the DOME Registry solution (Attafi *et al*. 2024), bringing together international stakeholders and experts from across Europe and beyond to collaborate on its establishment. The project delivered the foundational registry (Fig. 2), a robust platform for the controlled curation, management, and dissemination of supplementary AI method transparency reports to boost DOME compliance. The system allows researchers to directly create a structured, human-readable narrative detailing their publication’s AI methodology and providing at-a-glance transparency reporting metrics.

**Figure 2.**
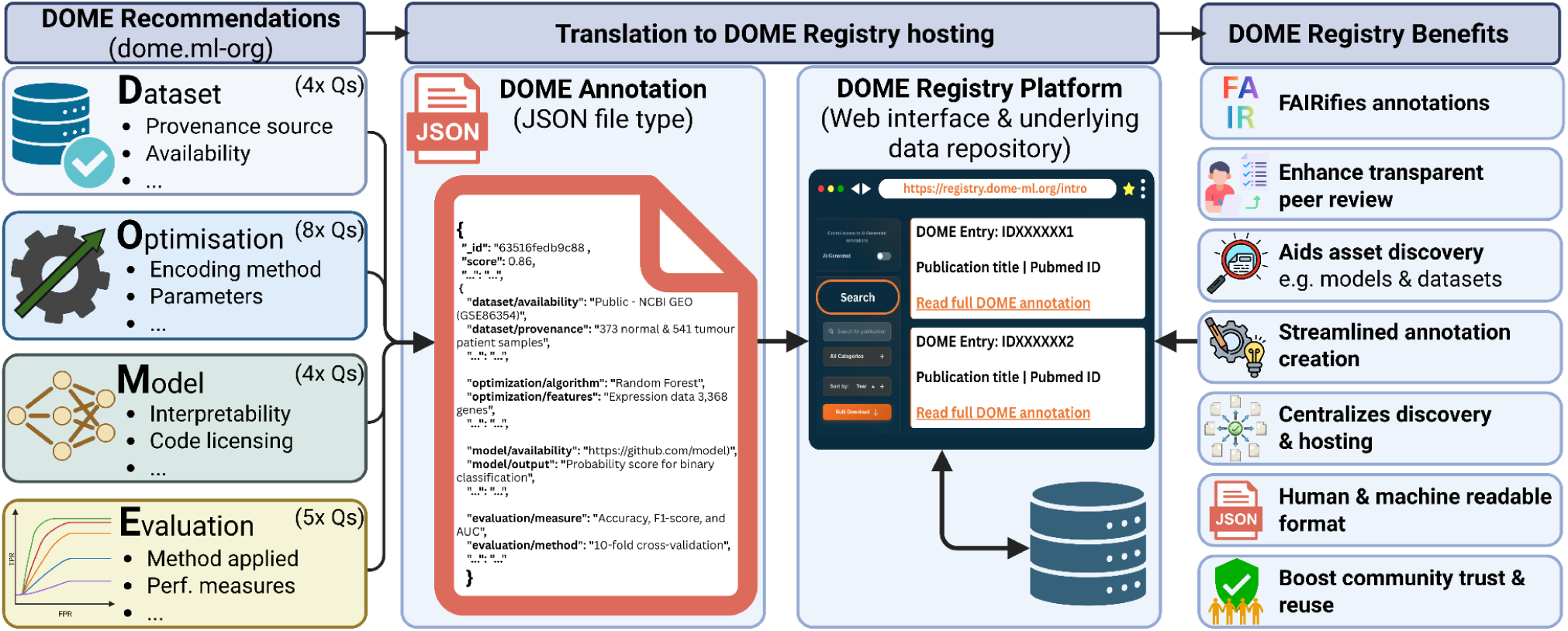
DOME Recommendations and Registry overview and benefits. Highlighted are the connections between the DOME Recommendations reporting guidelines and the DOME Registry platform with an example of a structured DOME transparency report in JSON format. The DOME Registry interface displays the human-readable narrative detailing the artificial intelligence methodology alongside compliance metrics supporting creating several benefits for the research community.

However, despite the best efforts of stakeholders across the scientific publishing chain to boost DOME adherence, challenges and barriers to adoption remained for widespread use of the system in the real-world. The reality of requiring authors to curate their own annotations in the DOME Registry presented a major overhead, with several caveats. For instance, the time required to report on a simple method is approximately one hour to properly curate, while complex methods can take up to three to four hours in total. At such a rate, achieving full coverage of the AI publication estimates shown in Fig. 1, assuming even 10-20% of the 619,980 results hold direct AI methods, this yields a crude estimate of 62,000 to 124,000 publications. At a rate of one hour per publication, it would take over 7 to 14 years of non-stop curation effort to cover the existing literature manually. This does not account for the continued increase in publications, making manual curation a Sisyphean task. Community curation incentives were employed, such as ORCID researcher profile (ORCID 2012) integrations to accredit and highlight DOME method curation via APICURON (Hatos *et al*. 2021), alongside training to help researchers easily annotate methods. Nevertheless, these attempts to address this manual incentive gap did not yield widespread adoption. Journal integration saw the most success in encouraging new annotations at the point of journal submission. However, as revealed in interviews with stakeholders from several journals, the publishing ecosystem is already slow and overburdened during review stages. Consequently, any additional strain on the system or time-consuming steps such as DOME reporting would not be viable for the average life sciences journal, despite the benefits it adds.

Considering the carrot-and-stick approach towards a solution for DOME compliance, it is difficult to find convincing, low-hanging fruit to ensure researchers adhere to DOME reporting or any other AI method reporting equivalents. If researchers can still secure funding, publish in journals, and sustain their careers without adhering to transparent method reporting, researcher culture will not adapt to integrate an additional step seen as a burden. Regarding the stick approach, as noted, adding DOME as a requirement in journal publication workflows through mandatory adoption did yield compliance and a degree of success, but the quality of annotations was observed to be variable, and in some cases rushed, resulting in minimal and lethargic annotation completion simply to satisfy the requirement.

The necessity for clear methods reporting is obvious from the reader’s perspective, but this is often not the case on the side of authors, who face intense pressure to publish. Additionally, for each case of journal adoption, the dedicated staffing required to operate the DOME Registry service would grow and necessitate further user support, as well as varied journal service-level agreements for access. This creates an intractable problem with major scalability hurdles, ultimately requiring a professionalized and for-profit service, which is not the goal for DOME. Furthermore, with the onslaught of mass-produced AI publication submissions and journals under strain, introducing additional requirements or steps, beyond a broad recommendation for compliance, is difficult to advocate for in an already overwhelmed system. Finally, the lack of incentives for curation, combined with it being viewed as ghost work not captured in the chain of academic assessment, acts as a further disincentive to engage in such method curation efforts (Davies 2025). Therefore, a novel solution was needed to address these major challenges (Table 1).

**Table 1.**
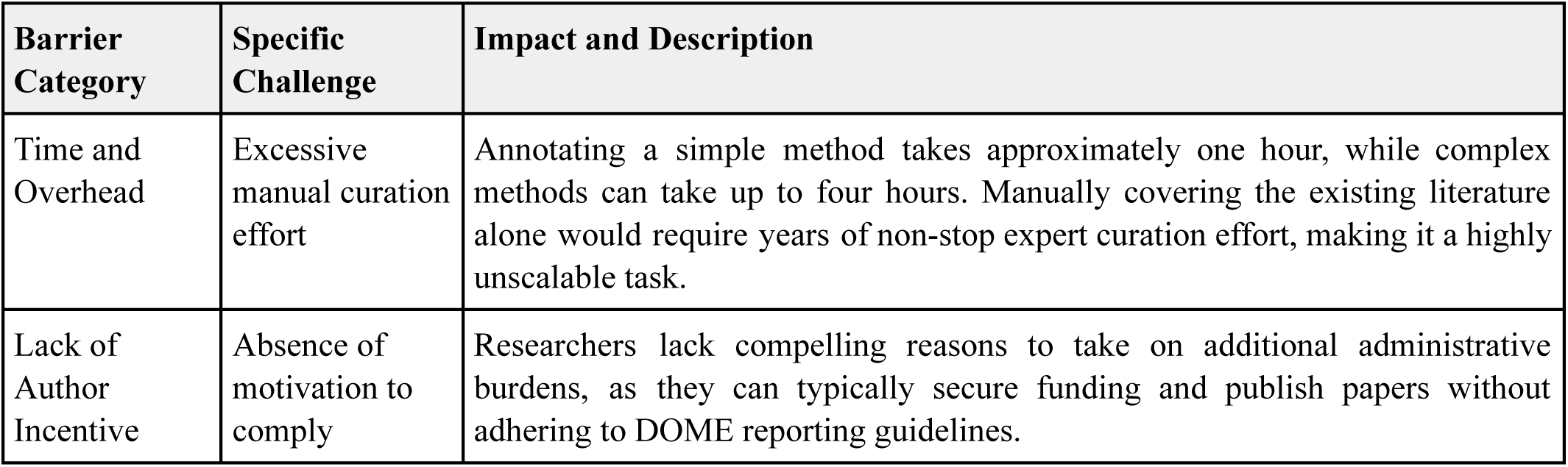
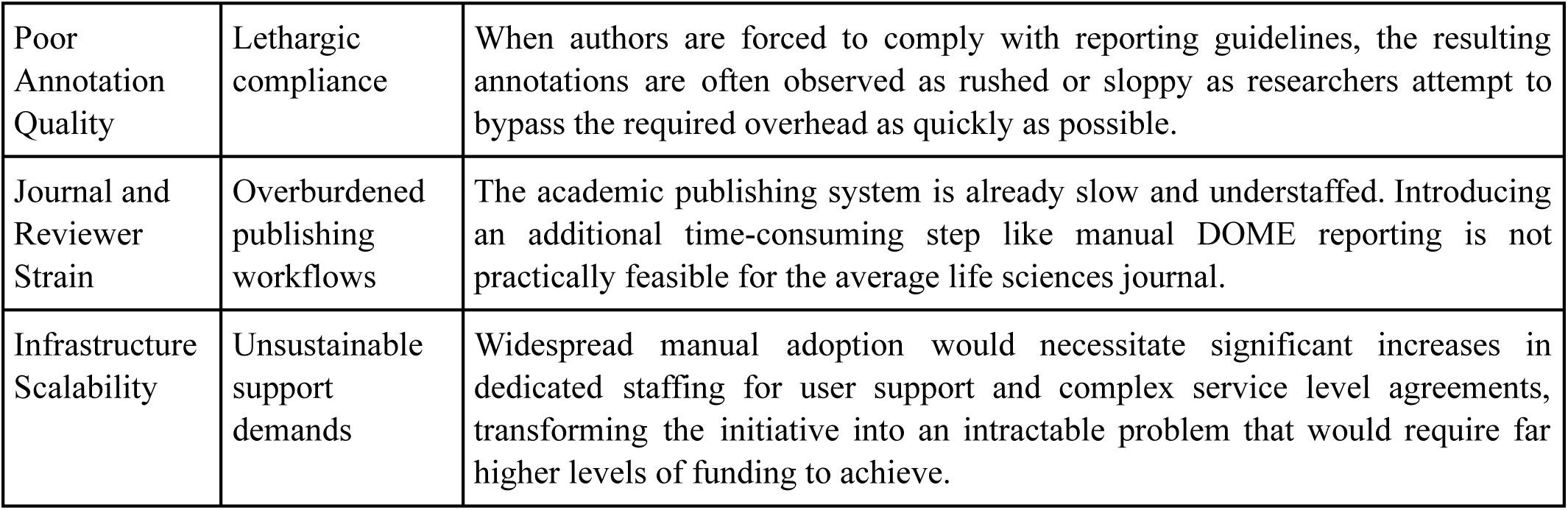
DOME Recommendations & Registry barriers to adoption and report creation.

| Barrier Category | Specific Challenge | Impact and Description |
| --- | --- | --- |
| Time and Overhead | Excessive manual curation effort | Annotating a simple method takes approximately one hour, while complex methods can take up to four hours. Manually covering the existing literature alone would require years of non-stop expert curation effort, making it a highly unscalable task. |
| Lack of Author Incentive | Absence of motivation to comply | Researchers lack compelling reasons to take on additional administrative burdens, as they can typically secure funding and publish papers without adhering to DOME reporting guidelines. |
| Poor Annotation Quality | Lethargic compliance | When authors are forced to comply with reporting guidelines, the resulting annotations are often observed as rushed or sloppy as researchers attempt to bypass the required overhead as quickly as possible. |
| Journal and Reviewer Strain | Overburdened publishing workflows | The academic publishing system is already slow and understaffed. Introducing an additional time-consuming step like manual DOME reporting is not practically feasible for the average life sciences journal. |
| Infrastructure Scalability | Unsustainable support demands | Widespread manual adoption would necessitate significant increases in dedicated staffing for user support and complex service level agreements, transforming the initiative into an intractable problem that would require far higher levels of funding to achieve. |

To bypass these systemic bottlenecks and overcome the unscalable nature of manual curation highlighted in Table 1, infrastructure providers in the Open Science and research infrastructure space are turning to automated curation. These stakeholders are seeing widespread success with the application of LLMs across areas such as RNA data resources (Green *et al*. 2026, The RNAcentral Consortium *et al*. 2026) and traditional knowledge bases for a diverse range of tasks (Wood *et al*. 2026). This success is further exemplified by a suite of related AI curation training, tools, and strategies such as those pioneered by Chris Mungall and collaborators, including AI agent-focused ontology management capacities for the Open Biological and Biomedical Ontologies (OBO) (Mungall 2025b), alongside the Monarch Initiative’s CurateGPT tool (Caufield *et al*. 2024) and the AI4curation community-focused resource (Mungall 2025a).

As a result of these community successes, the DOME initiative prioritized the development of a low-overhead AI-driven solution to significantly boost automated curation capabilities of AI transparency reports. By employing a human-in-the-loop approach, the goal of the system concept was to reduce the time involved in annotation, allowing researchers to simply review and finalize reports rather than writing them out manually. This would save time, minimize administrative burdens, and increase the attractiveness of DOME journal integration, thereby allowing the system to scale effectively and achieve better coverage of transparent AI methodologies. Therefore, the DOME Copilot was conceived as the next step in the evolution of the DOME framework, designed to address the scalability challenges of creating standardized reports to improve the transparency and reproducibility of AI methodology in the literature. It leverages the concept of AI for FAIR, whereby AI is used to increase the FAIRness of a resource through methods such as automated data curation. Accordingly, the DOME Copilot was designed with several key objectives: to be a lightweight, accessible, open-source, and reusable system; to maximize ease of use; and to facilitate human-in-the-loop final edits. Ultimately, this boosts the global coverage of AI methodology reporting, helping to restore trust and value in the corpus of published AI methods.

## Methods

### Datasets

The ground-truth human annotations were sourced from the DOME Registry (Attafi *et al*. 2024), and the benchmark dataset composed of manually curated DOME AI method reports. These entries provided the comparative baseline required to benchmark the performance of automated DOME Copilot outputs against human-annotated ground-truth. To download the corresponding source literature, first a Python script was developed to retrieve the full DOME Registry-hosted JavaScript Object Notation (JSON) entries via the registry’s application programming interface (API). The registry metadata included digital object identifiers (DOIs), which were then converted to their corresponding PubMed identifier (PMID) (FAIRsharing Team and Anderson 2023) and PubMed Central identifier (PMCID) (FAIRsharing Team 2023) using the National Center for Biotechnology Information (NCBI) E-utilities (Sayers 2009). From the initial pool of *n* = 258 DOME Registry manual annotations available, the pipeline resulted in a final set of *n* = 222 PMCIDs for which the full-text publications and associated supplementary materials were successfully retrieved in PDF format.

In addition to the human-curated benchmark dataset, two further datasets were generated. These datasets represent the DOME Copilot processing outputs across iterations: the initial prototype annotations and the final model annotations. These datasets consist of the DOME Copilot AI-generated transparency reports used for comparative evaluation against the human-annotated ground truth reports of the DOME Registry. These datasets facilitated the progressive assessment of the DOME Copilot’s performance between the model development iterations. The specific model processing, prompt refinement logic, and technical architectures related to these datasets are further detailed in the subsequent methodology sections.

### Models Selection and Justification

In alignment with the FAIR principles to support openness and reusability, open-source models were explicitly chosen for the DOME Copilot architecture. The Qwen3-Embedding-4B model (Zhang Y *et al*. 2025) was selected for text embedding generation due to its optimal balance of performance and efficiency. It achieved performance comparable to state-of-the-art models, ranking fifth on the Massive Text Embedding Benchmark (MTEB) (Muennighoff *et al*. 2022) leaderboard on Hugging Face at the time of writing, with significantly low latency. For the core annotation generation, Mistral-Small-3.1-24B-Instruct-2503 (Mistral 2025) was utilized. This intermediate-sized language model was chosen as it provides the capabilities required for complex text summarization and structuring while maintaining the inference speeds necessary for a scalable and responsive processing workflow.

This open-source strategy also addresses economic constraints, as recently escalating token costs for frontier commercial models (AI Business Review 2026) pose a long-term sustainability risk despite their performance advantages. The DOME Copilot system was designed to be flexible and can seamlessly integrate other models via APIs, as enabled by the Gradio (Abid *et al*. 2019) framework. This flexibility means that adoption by commercial publishers requiring the highest possible performance is feasible, even at a higher cost.

### Iterative Development and System Optimization

The development of the DOME Copilot involved key decisions and iterative adjustments to improve the outputs from the initial prototype to the final version. The fine-tuning of the prompts and model hyperparameters, such as the number of embedding nodes in the retriever and the PDF parser settings, was conducted using feedback from a benchmark validation subset. This subset consisted of *n* = 30 manual annotations from the benchmark dataset compared against the corresponding DOME Copilot prototype-generated set. In order to avoid data leakage in the reported results of the evaluation section, this prototype benchmarking dataset was kept separate from the final evaluation dataset. The remaining subset of *n* = 192 manual annotations was used for the semantic similarity analysis which used algorithmic metrics such as the BERTScore (Zhang T *et al*. 2019).

The majority of refinements implemented in the final system focused on improving the model processing prompts regarding expected style, formatting, and redundancy. Guided by observations from expert curators who identified inadequate patterns in the generation of annotation fields, measurable improvements were successfully implemented. For example, the succinctness of the final annotations was significantly improved by reducing verbosity. Multiple refinement iterations were applied across the three prompts (preprocessing, system, and section-specific) to support the detection of relevant DOME reporting information and the enforcement of structural constraints until the extraction of sufficient information and the desired composition of the generated annotations were achieved.

### User and Model Inference Workflow

The finalized architecture consisted of an LLM enabled with retrieval-augmented generation and structured prompt guidance. The submission workflow begins with a user-friendly Gradio interface where a researcher can upload a publication manuscript (PDF) along with optional supplementary materials. The inference workflow built with LlamaIndex (Liu 2022) is then triggered. First, a preprocessing step evaluates whether the PDF is relevant and contains AI methods. If it passes, the DOME Recommendations sections are iteratively generated; if not, it is rejected, refusing annotation.

For text extraction, the PDF contents are parsed with pypdf (Fenniak 2024). Docling (Auer *et al*. 2024) was considered as an alternative document content extractor as it theoretically offers advantages for structure preservation and features direct LlamaIndex integrations. However, in practice, it performed similarly to or worse than the more basic pypdf extractor while inducing significantly longer processing times. Therefore, pypdf was chosen as the default during the workflow, facilitating the best compromise between speed and accuracy.

For publication metadata retrieval (e.g., title, authors, journal, year), the Crossref API (FAIRsharing Team and Lister 2020) is used to directly retrieve accurate metadata based on user-provided DOIs. Where no metadata is retrievable, backup APIs trigger for alternative sources such as relevant preprint servers including bioRxiv (Cold Spring Harbor Laboratory 2013) and arXiv (Cornell University Library 1991). In cases where the optional DOI is not provided, the DOME Copilot extracts the information directly from the PDF contents. Inclusion of a DOI source supporting API metadata retrieval is preferred to ensure standardized, interoperable publication metadata for annotations intended for public hosting on the DOME Registry.

Once the text is extracted, the Qwen3-Embedding-4B model generates the embeddings from the text chunks, storing them in a local vector database integrated in LlamaIndex. To generate the full set of DOME annotation fields, the system iterates over the individual sections. Each query is composed of a system prompt and a section-specific prompt. The system prompt describes the context for the model, the expected style, and targeted instructions on how the model should react under predefined situations (such as outputting standard terminology indicating missing information when the model lacks context). The section-specific prompt consists of titles, a general description, and reference DOME questions the model is expected to answer, alongside formatting output instructions.

The section-specific prompt is embedded and compared with the vectors available in the local vector database. The top five most similar vectors are retrieved and their text chunks injected into the query as context. It was found that setting K=5 for the *top-K* similarity text retrieval was the best compromise to avoid missing important information and remain concise enough to avoid injecting irrelevant text noise.

This final prompt is then fed to the generative model (Mistral Small 3.1 24B Instruct 2503) using default hyperparameters (temperature=0.1). The response annotations are streamed in real time to the Gradio user interface for immediate progress visualization, taking approximately four minutes per run depending on the ingested document size. Upon completion, the annotation report can be downloaded directly as a structured JSON schema file for internal use, or edited and sent to the DOME Registry for public hosting. Future upgrades to the user interface are planned to close this operational loop, allowing users to review, edit, and directly submit finalized annotations entirely within the system.

### Deployment and Environmental Impact

Both the embedding and the generative models are deployed within the AI4EOSC (Heredia *et al*. 2025) platform. Specifically, they operate as part of the AI4EOSC LLM component running on two NVIDIA V100 for the generative model and one NVIDIA T4 GPU for the embedding model. To ensure full reproducibility and ease of redeployment, a Docker image is provided alongside the infrastructure code in the associated GitHub repository linked in the Code Availability section.

The total computational time required for the fine tuning and testing phases was 10 days, while the average inference time per manuscript is four minutes. The estimated carbon footprint and environmental impact of this compute, calculated using the MLCO₂ Machine Learning Emissions Calculator (Lacoste *et al*. 2019) was calculated at an upper bound estimate of 50 kg CO₂e emissions.

### Manual Performance Evaluation

To evaluate the performance of the DOME Copilot, a multi-staged approach involving both manual expert review and automated semantic scoring was implemented (Fig. 3). Initially, a subset of *n =* 30 human annotations from the primary dataset was selected to establish a comparative baseline against the DOME Copilot prototype outputs. This sample size represents a necessary compromise between achieving statistical significance and the intensive manual effort required for expert evaluation. The review focused on 630 individual field assessments (21 methodology fields across 30 publications), deliberately excluding publication metadata that the system can retrieve via definitive API calls.

**Figure 3.**
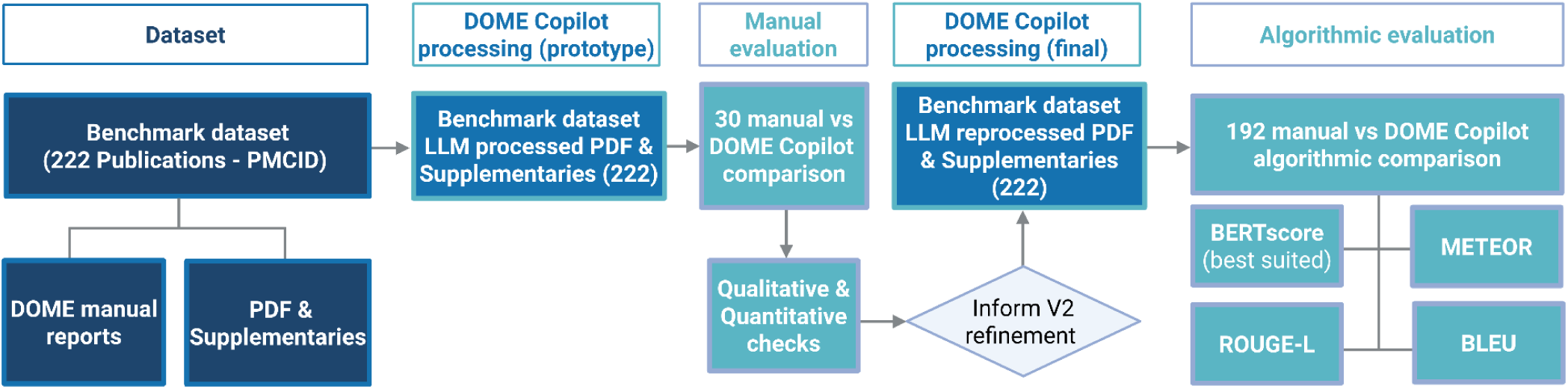
Overview of the DOME Copilot development workflow and evaluation methods. The primary dataset consisted of 222 publications identified by PMCIDs with existing manual DOME reports and associated full-text and supplementary PDFs. The full-text and supplementary PDFs were processed by the DOME Copilot prototype to trial automated generation via an LLM. Thirty of these automated reports were reviewed via manual comparison, leading to qualitative and quantitative information which informed the finalized DOME Copilot development. The final DOME Copilot was used to reprocess the dataset, and the annotations of the 192 held out from the first evaluation were used for secondary evaluation via algorithmic methods including BERTScore (Zhang T *et al*. 2019), ROUGE-L (Lin 2004), BLEU (Papineni *et al*. 2002), and METEOR (Banerjee and Lavie 2005).

An expert curator then performed iterative cross-referencing between source manuscripts and supplementary materials to grade annotations using a four-point quantitative rubric (Table 2) based on accuracy, structure, and comprehensiveness. This process was instrumental in identifying qualitative shortcomings in the prototype, such as excess verbosity, redundancy, and structural inconsistencies, which guided the prompt refinement and structural constraints for the final model. Furthermore, a gold-standard validation case was conducted using the AlphaFold 2 (Jumper *et al*. 2021) publication detailed in Supplementary Material 1, where a high-quality entry curated by a domain expert served as a rigorous reference to validate the final system’s ability to extract and summarize highly complex AI architectures.

**Table 2.** Quantitative evaluation rubric used during the manual review comparing DOME Copilot prototype outputs against the human baseline dataset. Details the rating and corresponding evaluation criteria.

| Rating | Evaluation criteria explanation |
| --- | --- |
| DOME Copilot prototype objectively better annotation | The generated output was more accurate, structured, or comprehensive than the human baseline, or successfully identified methodology data that the human annotator missed. |
| Human objectively better annotation | The human ground truth was superior, typically because the DOME Copilot failed to extract nuanced details, struggled with multi-model complexities, or produced overly verbose and irrelevant text. |
| Both high-quality annotations | Both the DOME Copilot and the human curator successfully and accurately extracted the required methodology information to a comparable, satisfactory standard. |
| Both low-quality annotations | Neither output was satisfactory. This typically occurred when the source publication lacked sufficient methodological detail, resulting in poor or absent extractions from both the model and the human. |

To calculate performance significance, the qualitative ratings from the rubric were encoded into numeric preference scores: +1 for a DOME Copilot win, -1 for a human baseline win, and 0 for any tied outcome. Using these encoded data, model performance was robustly evaluated via the SciPy library (Virtanen *et al*. 2020). Specifically, a global one-sample *t*-test and a binomial sign test were applied to assess the overall mean preference and win proportion, alongside per-field one-sample *t*-tests to evaluate significance within individual methodology categories. It is acknowledged that the per-field *t*-tests rely on highly discrete data; however, the sample size (*n* = 30) satisfies the traditional minimum threshold for the Central Limit Theorem, providing a practical and interpretable measure of mean preference. A summary of these statistical tests is detailed in Table 3.

**Table 3.** Statistical tests applied for comparative evaluation. Summary of the methodologies used to determine the statistical significance of the DOME Copilot prototype performance against the human ground truth baseline.

| Statistical test | Application and methodology | Package used |
| --- | --- | --- |
| Global one-sample *t*-test | Applied to test if the mean preference score across all valid observations is significantly different from 0 (where 0 represents a tie). | https://scipy.org/<br>(Virtanen *et al.* 2020)<br>(`scipy.stats.ttest_1samp`) |
| Binomial sign test | Applied to test if the proportion of DOME Copilot wins is significantly different from human wins, ignoring ties, under the null hypothesis that the probability of either winning is equal ($p = 0.5$). | https://scipy.org/<br>(Virtanen *et al.* 2020)<br>(`scipy.stats.binomtest`) |
| Per-field one-sample *t*-test | Applied to each DOME field individually to determine if the mean score for that specific methodology field is significantly different from 0. | https://scipy.org/<br>(Virtanen *et al.* 2020)<br>(`scipy.stats.ttest_1samp`) |

### Semantic Scoring Performance

The broader performance of the final DOME Copilot was assessed through automated similarity metrics across the remaining *n* = 192 entries of the benchmark dataset. These entries were strictly withheld from the initial refinement process to ensure an independent evaluation and avoid data leakage. BERTScore was utilized as the primary metric for this analysis, as it leverages contextualized token embeddings from a pre-trained transformer backbone (microsoft/deberta-v3-base) (He, Gao, and Chen 2021) to compute pairwise cosine similarities. This semantic approach is specifically suited for comparing generative outputs against diverse, prose-based human ground-truth annotations, as it rewards contextual similarity rather than exact lexical overlap. While BERTScore served as the best test for comparative evaluation, other lexical and semi-semantic metrics, including ROUGE-L (Lin 2004), BLEU (Papineni *et al*. 2002), and METEOR (Banerjee and Lavie 2005) were also calculated to provide a comprehensive overview of textual relationships and structural coherence (detailed in Supplementary Material 2). This methodology allowed for a robust assessment of how the structured, succinct annotations generated by the final DOME Copilot align with the semantic intent of the original human-curated registry entries.

## Results

### Finalized DOME Copilot System Architecture

The final DOME Copilot was developed to provide a seamless and user-friendly experience for automated transparency reporting (Fig. 4). A video demonstration of the system in action is available online and linked in the Code Availability section. The system workflow is initiated when a user submits the main PDF of an AI methods manuscript, alongside an optional supplementary PDF or a DOI, via the Gradio user interface. Following the upload, text is extracted from the PDFs using pypdf. Where an optional DOI is provided, the publication metadata is extracted from authoritative external literature registries via APIs. During the subsequent processing phase, the primary workflow is orchestrated by LlamaIndex. The extracted text is first passed to the Qwen3-Embedding-4B model to generate vector embeddings. These vectors are then ready for LLM ingestion to process the DOME question set prompts using Mistral-Small-3.1-24B-Instruct-2503. Finally, the generated DOME report is returned via the Gradio user interface. The structured JSON report format is downloadable for further edits and can subsequently be uploaded to the DOME Registry for public sharing and hosting in a FAIR manner.

**Figure 4.**
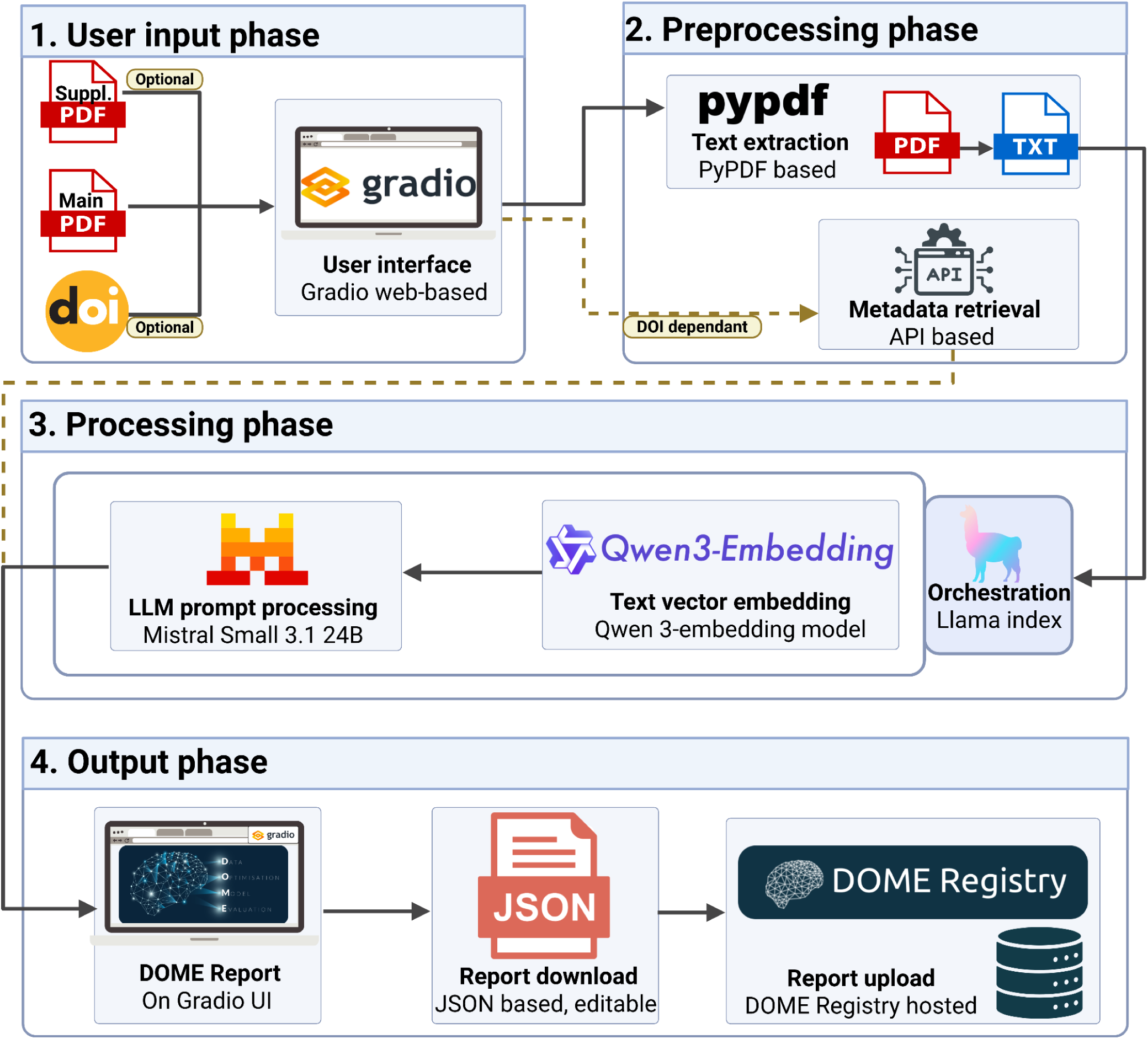
DOME Copilot system workflow overview. Schematic breakdown of the finalized DOME Copilot’s automated transparency reporting pipeline. The workflow progresses sequentially from user input of manuscript PDFs and optional DOIs via the Gradio interface, through raw text extraction using pypdf and API-driven metadata retrieval. LlamaIndex orchestrates the core annotation framework, utilizing Qwen3-Embedding-4B for text vectorization and a *top-k* similarity search (*k* = 5) to populate category-specific prompt templates. Final structured reports are synthesized by the Mistral-Small 3.1 24B generative model and streamed back to the user interface as a downloadable JSON file ready for human-in-the-loop validation and public DOME Registry integration.

### Qualitative Findings of the Manual Review

The qualitative observations gathered during the manual comparison of the DOME Copilot prototype outputs against the manual annotations were used to optimize the prompts and refine the output structure of the final DOME Copilot model to address several observed shortcomings are captured in Table 4.

**Table 4.** Qualitative shortcomings identified in DOME Copilot prototype. Summary of the primary issues observed during the manual expert review of the initial AI-generated outputs, which subsequently guided the prompt refinement and structural constraints implemented for the final model.

| Identified shortcoming | Observation |
| --- | --- |
| Excess verbosity | The model often provided overextended responses. |
| Redundancy | Outputs frequently contained redundant, albeit seemingly useful, information rather than stating when data was simply unavailable across DOME fields. |
| Not applicable misuse | The model incorrectly used "Not Applicable," whereas all DOME fields should be considered applicable and require a definitive answer or a clear statement of missing information. |
| Poor structure | A need was identified to enforce better structuring of entries and to more strictly extract URLs in a clearly formatted manner rather than embedding them in long text blocks. |
| Inconsistency | The model struggled with multi-question fields, often addressing only one aspect and varying its approach inconsistently. |
| Multi-model cases | The model had difficulty handling publications where more than one AI model was developed. |

### Quantitative Findings of the Manual Review

In addition to the qualitative observations, the manual review process enabled a quantifiable ranking evaluation of the final DOME Copilot annotations versus the corresponding human set across the 30 publications. The comparative ratings were made by expert curator judgement using the following four-point evaluation rubric described in Table 2. Applying this rubric to the benchmark validation subset provided a clear assessment of the initial DOME Copilot prototype model performance. This systematic scoring highlighted the specific DOME fields where the prototype model struggled most, directly informing the targeted prompt engineering required for the final version. Despite these identified areas for improvement, the aggregate field evaluations demonstrated that the initial AI extractions were already highly competitive. When isolating the core methodology fields, the prototype model objectively outperformed or tied with the human ground truth in the vast majority of instances (Fig. 5).

**Figure 5.**
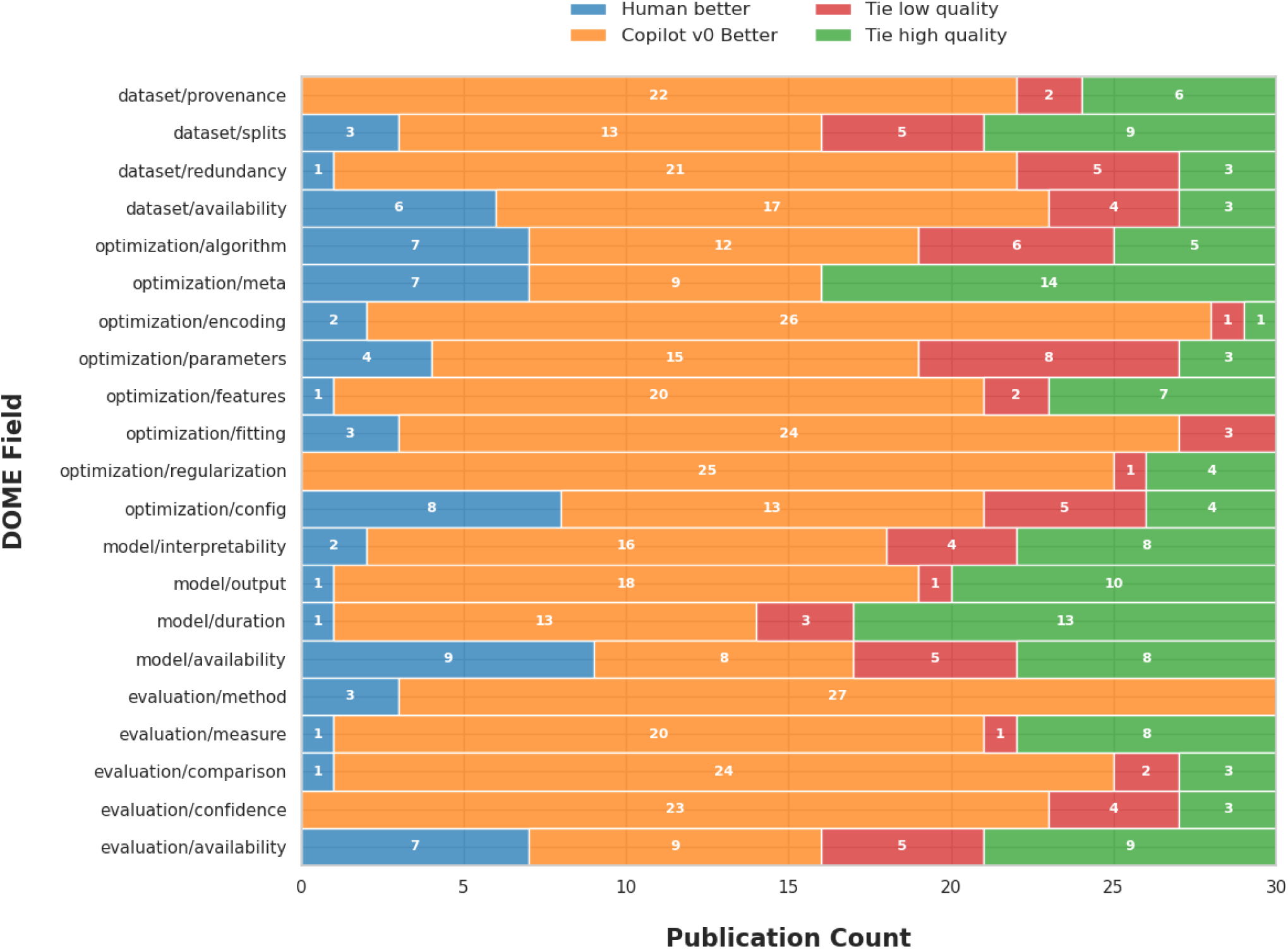
Comparative performance evaluation of DOME Copilot prototype versus manual annotations. Results from the manual review of the benchmark validation subset of 30 publications across 630 DOME fields, comparing the initial AI-generated outputs against the corresponding ground-truth manual annotations. The 21 stacked bars represent each individual DOME methodology reporting field, with four distinct colours representing the distribution of rubric classifications. This evaluation was utilized to identify specific areas of weakness that required the most improvement, directly guiding the prompt and structural refinements for the final model.

### Statistical Findings of the Manual Review

These field-level manual review evaluations allowed for calculating the following statistical findings comparing the DOME Copilot prototype performance against the human ground-truth. Analysing these computed tests demonstrated significant performance trends that actively guided focused prompt refinement to address model weaknesses and streamline outputs for the final release. All data were drawn directly from the custom manual review interface CSV-logged outputs by the expert curator (see Data Availability).

### Global Overview and Statistical Significance

The manual review process yielded 630 valid field-level observations (after excluding irrelevant publication metadata). An analysis of the global performance demonstrated a statistically significant preference for the DOME Copilot AI-generated outputs over the human ground truth. The DOME Copilot prototype won 59.5% of the evaluations compared to the human annotators’ 10.6%, producing an overall mean preference score of 0.489 (Table 5). Both the global one-sample *t*-test (favouring the DOME Copilot prototype) and the exact binomial sign test confirmed that this performance advantage is statistically significant (*p* = 5.27 x 10^-53^), conclusively rejecting the null hypothesis.

**Table 5.** Global statistical overview and significance of DOME Copilot prototype versus human manual annotations.

| Metric | Result |
| --- | --- |
| Total field observations | 630 |
| DOME Copilot prototype better (Wins) | 375 (59.5%) |
| Human better (Wins) | 67 (10.6%) |
| Ties (Total) | 188 (29.8%) |
| *High-quality tie* | *121 (19.2%)* |
| *Low-quality tie* | *67 (10.6%)* |
| Mean preference score | 0.489 |
| Global one-sample $t$-test (Two-sided test against pop. mean = 0) | T-statistic: 18.028<br>P-value: $6.95 \times 10^{-59}$<br>(Significant difference favouring DOME Copilot) |
| Binomial sign test (Exact test) | P-value: $5.27 \times 10^{-53}$<br>(Reject null hypothesis $H_0$ where $H_0$: $p=0.5$ (excluding ties). The win rate is not 50/50.) |

### Per-Field Analysis

To isolate the specific methodology areas driving this global significance, a per-field analysis was conducted using individual one-sample *t*-tests (Table 6). The DOME Copilot demonstrated a statistically significant advantage across the majority of the extraction tasks, particularly in the DOME fields related to optimization, encoding, and dataset provenance.

**Table 6.** DOME field level statistical analysis. Fields are ordered descending by mean preference score (representing the DOME Copilot’s advantage). The *P*-value column displays the result of a one-sample *t*-test for that specific field versus 0 (a tie). An asterisk (*) indicates statistical significance (*p* < 0.05).

| DOME field | Mean score | DOME Copilot wins | Human wins | Ties | $P$-value |
| --- | --- | --- | --- | --- | --- |
| optimization/regularization | 0.833 | 25 | 0 | 5 | $8.3 \times 10^{-13}$ * |
| evaluation/method | 0.800 | 27 | 3 | 0 | $6.6 \times 10^{-08}$ * |
| optimization/encoding | 0.800 | 26 | 2 | 2 | $9.0 \times 10^{-09}$ * |
| evaluation/comparison | 0.767 | 24 | 1 | 5 | $3.5 \times 10^{-09}$ * |
| evaluation/confidence | 0.767 | 23 | 0 | 7 | $1.1 \times 10^{-10}$ * |
| dataset/provenance | 0.733 | 22 | 0 | 8 | $8.1 \times 10^{-10}$ * |
| optimization/fitting | 0.700 | 24 | 3 | 3 | $2.2 \times 10^{-06}$ * |
| dataset/redundancy | 0.667 | 21 | 1 | 8 | $2.5 \times 10^{-07} *$ |
| optimization/features | 0.633 | 20 | 1 | 9 | $8.3 \times 10^{-07} *$ |
| evaluation/measure | 0.633 | 20 | 1 | 9 | $8.3 \times 10^{-07} *$ |
| model/output | 0.567 | 18 | 1 | 11 | $7.0 \times 10^{-06} *$ |
| model/interpretability | 0.467 | 16 | 2 | 12 | $3.4 \times 10^{-04} *$ |
| model/duration | 0.400 | 13 | 1 | 16 | $5.4 \times 10^{-04} *$ |
| optimization/parameters | 0.367 | 15 | 4 | 11 | $9.1 \times 10^{-03} *$ |
| dataset/availability | 0.367 | 17 | 6 | 7 | $1.9 \times 10^{-02} *$ |
| dataset/splits | 0.333 | 13 | 3 | 14 | $9.9 \times 10^{-03} *$ |
| optimization/algorithm | 0.167 | 12 | 7 | 11 | $2.6 \times 10^{-01}$ |
| optimization/config | 0.167 | 13 | 8 | 9 | $2.8 \times 10^{-01}$ |
| evaluation/availability | 0.067 | 9 | 7 | 14 | $6.3 \times 10^{-01}$ |
| optimization/meta | 0.067 | 9 | 7 | 14 | $6.3 \times 10^{-01}$ |
| model/availability | -0.033 | 8 | 9 | 13 | $8.1 \times 10^{-01}$ |

### DOME Copilot evaluation results (BERTScore: *n* = 192)

Performance in generating relevant annotations was evaluated through the calculation of BERTScore metrics. This provided a numeric measure of the contextual and semantic similarity between *n* = 192 entries of the overall human annotation benchmark dataset (*n* = 222) against those generated by the final DOME Copilot. The *n* = 30 benchmark dataset entries already used for the DOME Copilot refinement process during model development were excluded during BERTScore evaluation to avoid data leakage. The more classical evaluation metrics that were trialled are available in Supplementary Material 2, but were found to be unsuited to the evaluation task given their limitations in capturing contextual similarity and applicability, thereby reinforcing the validity of semantic-based evaluation methods.

The BERTScore results assessing the contextual similarity of the final DOME Copilot-generated annotations against human annotations were found to have upper and lower quartiles primarily between 0.35 and 0.50 (Fig. 6). A BERTScore of 1 is representative of an identical semantic match, and these results indicate stable semantic similarity between DOME Copilot-generated annotations and the corresponding human-curated entries. A limitation of BERTScore as a measure of the DOME Copilot’s real performance is that the ground-truth human annotations are highly diverse, originating from many individual annotators, generally prose-based in style, and vary in quality. For most DOME fields, scores cluster within a relatively narrow range. Minor variability is observed between individual questions, particularly within the Model category, where some fields exhibit a slightly wider spread of scores. In contrast, the Dataset, Optimization and Evaluation categories display more compact distributions, suggesting more uniform performance across these fields.

**Figure 6.**
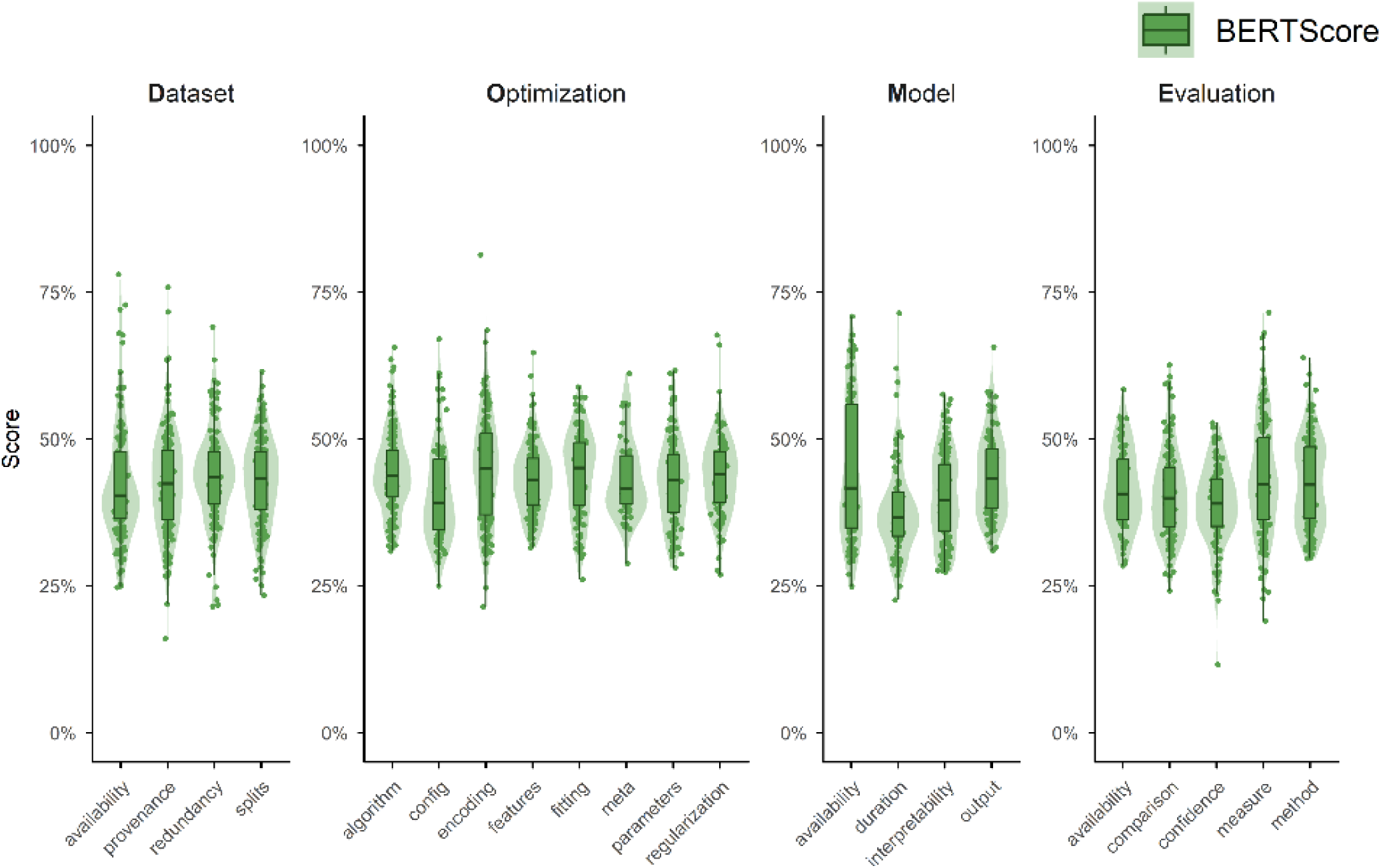
BERTScore semantic similarity evaluation of final DOME Copilot against manual annotations. The semantic similarity scores between the automated outputs and the ground-truth manual annotations for the holdout set of 192 publications. The results are grouped by the four core DOME categories (Data, Optimization, Model, and Evaluation) and span the 21 individual methodology fields. Boxplots illustrate the distribution of scores, with individual dots plotted outside the interquartile ranges to highlight extreme outliers.

## Discussion

### Strengths and Limitations of the Manual Validation of the Prototype Architecture

The manual evaluation of the prototype by the expert curator ranked that the DOME Copilot prototype was already capable of outperforming benchmark human annotated DOME reports by providing superior responses in complex topic areas where a human may have lacked specific domain expertise. In the instances where the DOME Copilot was observed to fail, these insights were directly leveraged for the final system refinements. This initial validation established a strong foundational performance baseline and identified the inherent limitations of manual curation even against the prototype model. This reflected the constraints such as human curator expertise, effort, and the incentive to perform annotations which impacted annotation quality – issues which the DOME Copilot prototype system successfully circumvented.

The primary limitation of the related statistical findings is the narrow sample size of the benchmark validation dataset. While the subset of *n* = 30 publications was carefully controlled to ensure balanced representation and satisfied the Central Limit Theorem for testing, an expanded review would yield greater statistical power. A broader sampling would better capture the rare edge cases and structural nuances present in highly complex AI methodologies. An additional limitation is the reliance on a single expert curator for the comparative grading. The inclusion of an independent co-curator would have enabled formal inter-rater reliability analysis to resolve subjective scoring variances. However, the prohibitive manual effort involved made this unfeasible at the time of study, as each individual review required up to two hours of intensive cross-referencing by a capable expert.

### Semantic Performance and Field-Level Interpretation of the Final Architecture

The algorithmic evaluation of the final DOME Copilot demonstrated moderate semantic similarity between the generated and manually curated annotations, with BERTScore yielding satisfactory performance metrics despite the limitations in direct comparison between human-curated and model-generated reports. A direct lexical match is not expected due to the highly diverse and prose-based style of the ground-truth human annotations. Consequently, other overlap-based lexical evaluation tests like ROUGE-L, BLEU, and METEOR (Supplementary Material 2) were found to be ill-suited to capturing this contextual similarity. The BERTScore distributions indicate a strong capability of the model and prompts to extract and carefully structure relevant methodologies rather than recreating highly variable human-like inputs.

At the DOME reporting field level, the DOME Copilot prototype primarily struggled with identifying URLs, a limitation exacerbated when such links were absent from the core text. This was observed in fields requiring explicit URL returns, such as model availability, evaluation availability, and optimization configuration. Human annotators demonstrated superior proficiency in locating these details by searching external sources like GitHub repositories and related web interfaces. Licensing was similarly affected; while the DOME Copilot could often retrieve a generic URL, human annotators were consistently better at explicitly identifying specific licensing details. Additionally, determining whether a model functions as a meta-predictor proved challenging for the DOME Copilot, often producing excessive text instead of a concise binary response. The optimization algorithm field also suffered from the complexities of multi-model publications, resulting in outputs that lacked the targeted succinctness achieved by manual annotations. Despite these specific challenges, the DOME Copilot demonstrated significant strengths in extracting complex, granular technical parameters. This capability was highlighted during the validation using the highly complex AlphaFold 2 methodology (Supplementary Material 1).

### Value of the DOME Copilot System and Current Use Cases

The system demonstrates notable value by offering a scalable, reliable solution capable of providing comprehensive reporting coverage across the 21 DOME fields. By removing the manual annotation bottleneck, it enables journal and author adoption in publishing workflows while helping researchers understand the full corpus of AI methods. Using modest compute and token costs, it provides a sustainable, and reusable approach contrasting favorably with the expensive overhead associated with commercial frontier models.

The DOME Copilot is capable of supporting three core use cases. First, as a self-checking aid, it assists developers in rapidly identifying methodological gaps requiring more detailed descriptions. Second, as a publishing workflow aid, it facilitates the submission process by automatically drafting a structured DOME methods transparency report. This provides publishers and reviewers with standardized method descriptions, reducing the administrative overhead of mandating reporting guidelines. Third, it enables scalable annotation coverage for the DOME Registry, supporting the reuse of archival and newly published methods. For the first two applications, users must carefully review and refine the output, maintaining human-in-the-loop workflows to correct potential errors while greatly reducing curation time. For the third application involving bulk archival processing, manual validation would be practically unfeasible. Therefore, any prospective processing of archival bulk publications will be transparently labelled in the DOME Registry to explicitly inform users that no manual refinement has occurred. By supporting these key use cases, the system significantly boosts methodological transparency and reproducibility.

### Future Directions

The DOME Copilot system will continue to improve performance as more advanced open-source models, refined embeddings, and wider context windows become available for integration into its flexible infrastructure. Other performance improvements will investigate the addition of agentic search and retrieval-augmented generation functionalities (Lewis *et al*. 2020) to explore URL cross-linked resources beyond the primary publication, such as external data and code repositories where additional information is likely to be held.

As AI systems evolve at a rapid pace, traditional and highly explainable frameworks like the DOME Recommendations are needed more than ever to avoid the black-box oracle paradox of AI in research (Wei *et al*. 2025). Therefore, embedding the DOME Copilot as a core component of the interconnected life science research infrastructure ecosystem will be undertaken, leveraging further collaboration across key Open Science services, including integrations with EBI Search (Pearce et al., 2025) and Europe PubMed Central to enhance discoverability at publication source. Additionally, tighter resource coupling with the DOME Registry interface will streamline user navigation, while alternative deployment options will be supported through the AI4EOSC platform (Heredia *et al*. 2025), including the potential for integration at national European Open Science Cloud (EOSC) Nodes (EOSC Association 2025).

Another major next step will look towards continued DOME journal integration and deepening engagement with academic publishers. By actively demonstrating how the tool mitigates the original administrative burden of DOME reporting for editors and reviewers, this is hoped to encourage widespread adoption within standard journal publication workflows. The DOME Copilot’s modular architecture will allow key stakeholders, such as publishers, to utilize their own compute infrastructure to establish private deployments. These localized deployments could then seamlessly connect to the wider DOME Registry via APIs for controlled report releases supplementing their publications.

## Conclusion

The DOME Copilot addresses the critical manual curation bottleneck in generating standardized AI method disclosures, offering a scalable and robust framework for researchers, publishers, and funding bodies to enhance the reusability and reproducibility of scientific literature applying AI methodologies. By automating the extraction and structuring of granular methodological information directly from manuscript PDFs, the system provides a powerful proof-of-concept for mitigating the scalability limitations inherent in manual compliance reporting. Looking ahead, the initiative aims to expand its operational availability by exploring deployment within European Union AI Factories, leveraging the AI4EOSC platform, and piloting publisher-led integrations. Through flexible API architectures and the ability for institutional stakeholders to leverage their own computational infrastructure, such as commercial cloud environments, the DOME Copilot is poised for widespread adoption into the global research infrastructure ecosystem, aiding methodological transparency of the expanding corpus of AI-driven life science research.

## Data and Code availability

**Zenodo**: https://doi.org/10.5281/zenodo.18615687

**GitHub DOME Copilot Data Preparation, Graphing & Analysis**: https://github.com/gavinf97/DOME-Copilot-Data-Analysis

**GitHub DOME Copilot Infrastructure**: https://github.com/IFCA-Advanced-Computing/dome-copilot

**Gradio DOME Copilot Access:** https://dome-copilot.ifca.es/ (reviewer login access provided during journal submission and also general public availability upon request to)

**DOME Copilot Demonstrator Video:** https://www.youtube.com/watch?v=iIMjrchceGg

## Acknowledgements

This work has been supported by: ELIXIR, the European infrastructure for life science data, through the ELIXIR projects 2023-MLstandards [F.P., S.C.E.T] and 2024-TECHNOLOGY-Data [M.H., M.J.]; EMBL Core funding [H.H., M.H., M.J, M.P.]; Consejería de Educación, Formación profesional and Universidades del Gobierno de Cantabria via the ‘Actividad estructural para el desarrollo de la investigación del Instituto de Física de Cantabria’ project [A.L.G.]. Additional funding from the European Union through NextGenerationEU PNRR project ELIXIRxNextGenIT (grant agreement no. IR0000010) [S.C.E.T.]; Horizon Europe projects AI4EOSC (grant agreement no. 101058593) [A.L.G.], EVERSE (grant agreement no. 101129744) [S.C.E.T., F.P.] and ELIXIR STEERS (grant agreement no. 101131096) [S.C.E.T., H.H., F.P.]. Views and opinions expressed are those of the author(s) only and do not necessarily reflect those of the European Union or the European Research Executive Agency. Neither the European Union nor the granting authority can be held responsible.

## Author contributions

All authors contributed to the DOME Copilot project work and subsequent discussion as well as the initial draft of this manuscript. G.F., O.A.A, S.C.F and I.H. as co-first authors wrote the initial draft with the help of all co-authors. All authors edited and refined the final manuscript. A.L.G. F.P. and S.C.E.T., the co-corresponding authors, initiated and coordinated the project.

## Correspondence

All correspondence should be addressed to Alvaro López García, Fotis Psomopoulos, or Silvio C. E. Tosatto.

