## Supplementary Material 1 for "DOME Copilot: A resource to automate transparent reporting of artificial intelligence methods"

### Supplementary Material 1.
### DOME Copilot vs Human: AlphaFold 2 Complex Manual Review Comparison

As a final evaluation of the refined DOME Copilot system, the flagship use case of the AlphaFold 2 publication (Jumper *et al.* 2021) was selected to undergo the same manual review methodology previously applied to the initial benchmark dataset subset ($n$ = 30) used for prototype refinement. The AlphaFold 2 specific manuscript represents a complex flagship AI method, spanning multiple training datasets and intricate architectural complexities. The final methodology consists of numerous distinct AI methodology considerations that must be accurately described and structured within the DOME reporting fields.

Comparing the DOME Registry's expertly curated human ground truth for this publication against the outputs generated by the finalized DOME Copilot provides a test of the model's extraction and summarization capabilities under highly complex reporting conditions. The results of this manual review, detailing the per DOME field comparative performance, are directly presented in Supplementary Figure 1.1.

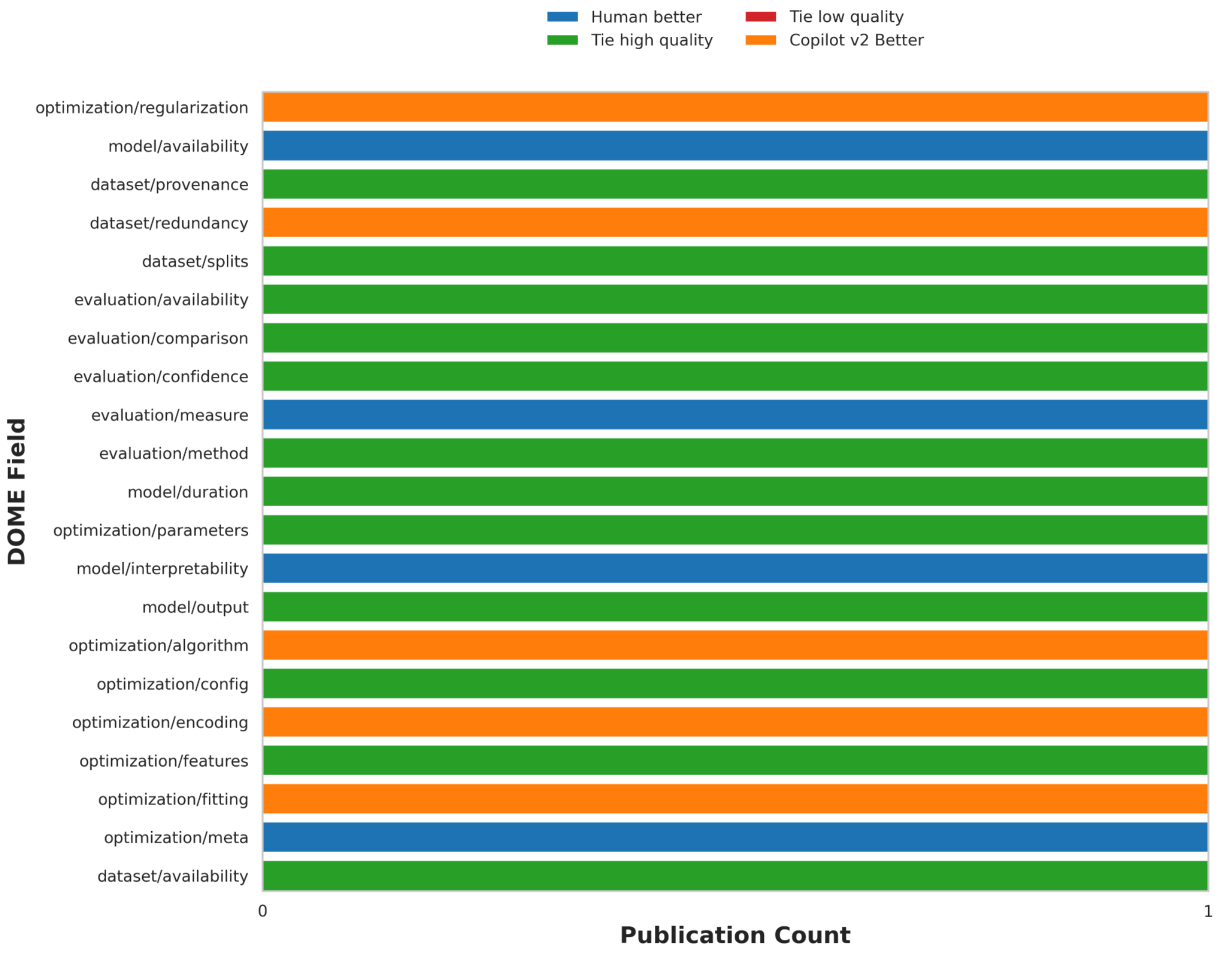


**Supplementary Figure 1.1. Comparative performance evaluation of the final DOME Copilot versus human-curated annotations for the AlphaFold 2 manuscript.** Results from the manual review of the AlphaFold 2 publication, comparing the final DOME Copilot AI-generated outputs against the corresponding expert ground-truth manual annotations across all required DOME methodology reporting fields.

The overall quantitative distribution of these comparative ratings is visualized in Supplementary Figure 1.2 to highlight the broader performance trends of the final DOME Copilot model outputs versus the human ground-truth annotation. Commentary was additionally collected during the comparisons during this manual review for qualitative observation collection between the two annotations. The exact dataset and generated outputs for this specific comparison are made available in the associated code repository.

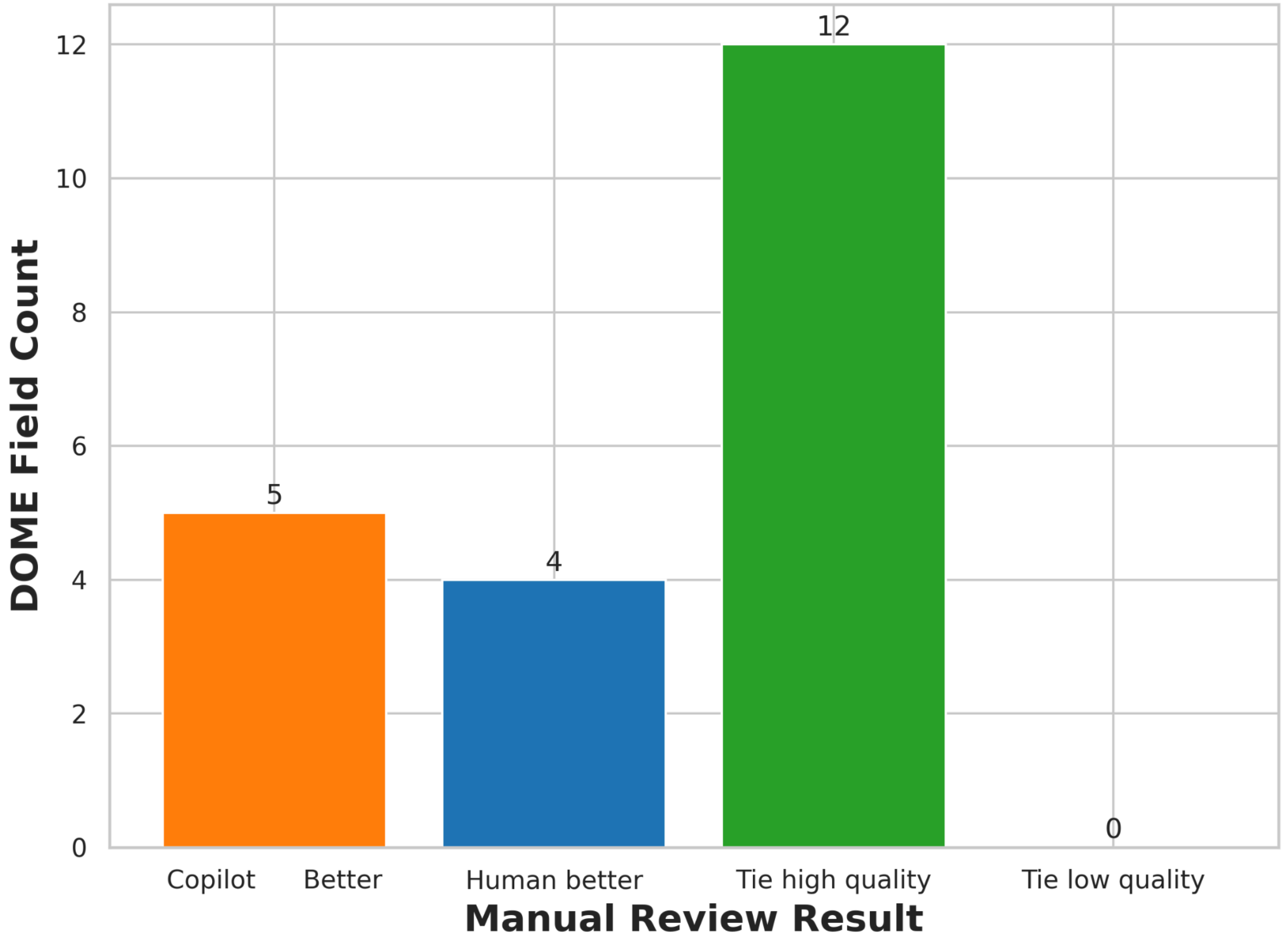


**Supplementary Figure 1.2. Quantitative evaluation distribution of final DOME Copilot versus human-curated annotations for the AlphaFold 2 manuscript.** The distribution of comparative ratings across the AlphaFold 2 DOME methodology fields, scored using the established four-point rubric previously used during the prototype manual review. The results highlight the proportion of fields where the final DOME Copilot outperformed the human ground truth, where the human baseline was superior, and where both outputs yielded ties.

**Overview of AlphaFold 2 Benchmarking Results**

The manual review highlights that the final DOME Copilot is effective at handling dense technical AI methodology data, though subtle nuances in model architecture, such as the degree of interpretability and external resource URL linking, remain areas where human expertise holds an advantage.

**Data Reporting Fields**

In evaluating dataset inputs, both the human curator and the final DOME Copilot successfully mapped the diverse genomic and structural frameworks utilized during model training. The automated pipeline effectively isolated structural data provenance links, capturing repositories like the Protein Data Bank (Berman *et al.* 2000) and UniRef90 (Suzek et al. 2015). Furthermore, the DOME Copilot provided a deep breakdown of the sequence identity clustering thresholds and the specialized filtering sequences used to maintain test set independence, earning a superior rating for its handling of the dataset redundancy field.

**Optimization Reporting Fields**

When analysing optimization metrics, the final DOME Copilot demonstrated a distinct capacity for capturing granular algorithmic variables, though it occasionally overlooked macro-level architectural classifications. The automated framework generated a detailed description of data encoding mechanisms, explicitly registering the specific one-hot encoding dimensions for primary amino acid sequences and multi-sequence alignment blocks. It likewise tracked essential regularization and training stabilization configurations, such as spatial dropout thresholds. Conversely, the human curator displayed a stronger grasp of abstract system elements, successfully documenting the unique structural patterns of the Evoformer layers and graph neural networks, whereas the DOME Copilot miscategorized the complex pipeline as a meta-predictor

**Model Reporting Fields**

For model-specific criteria, the evaluation revealed a high-quality tie across the majority of execution and hardware categories. Both curation pathways accurately documented the heavy resource footprints of the framework, logging the multi-week hardware training runs and exact inference latency speeds. The human curator achieved a higher mark for interpretability by thoroughly tracing the authors' explicit explainability frameworks, while the final DOME Copilot initially defaulted to treating the system as a closed black-box architecture, missing the explicit manuscript segments dedicated to ablation studies and network visualization efforts. On resource accessibility, the human expert pinpointed the precise URL path to the source repository, whereas the DOME Copilot resolved its extraction to the higher-level organization subdirectory.

**Evaluation Reporting Fields**

In the assessment of validation benchmarks, the final DOME Copilot proved highly competitive when isolating core testing protocols, though it captured a more consolidated profile of secondary performance metrics than the human expert. Both approaches successfully identified the primary validation baseline, noting the integration of the independent Critical Assessment of Structure Prediction 14 (CASP14) (Kryshtafovych *et al.* 2021) dataset. The automated pipeline accurately extracted primary spatial precision coordinates, including predicted Local Distance Difference Test (pLDDT) and Template Modeling score (TM-score). The human curator, however, provided a more exhaustive matrix of CASP-specific parameters, cataloguing the area under the receiver operating characteristic curve (AUC-ROC) for residue contact predictions alongside secondary correlation coefficients. Both correctly identified the reporting of 95% confidence intervals, confirming the mathematical reliability of the underlying structural predictions.

**References (Supplementary Material 1)**


Berman HM, Westbrook J, Feng Z et al. The Protein Data Bank. *Nucleic Acids Res* 2000;28:235–42. https://doi.org/10.1093/nar/28.1.235

Jumper J, Evans R, Pritzel A et al. Highly accurate protein structure prediction with AlphaFold. *Nature* 2021;596:583–9. https://doi.org/10.1038/s41586-021-03819-2

Kryshtafovych A, Schwede T, Topf M et al. Critical assessment of methods of protein structure prediction (CASP)—round XIV. *Proteins* 2021;89:1607–17. https://doi.org/10.1002/prot.26237

Suzek BE, Wang Y, Huang H et al. UniRef clusters: a comprehensive and scalable alternative for improving sequence similarity searches. *Bioinformatics* 2015;31:926–32. https://doi.org/10.1093/bioinformatics/btu739
