## Supplementary Material 2 for "DOME Copilot: A resource to automate transparent reporting of artificial intelligence methods"

### Supplementary Material 2.
### DOME Copilot vs Human: Similarity Metric Evaluations

To evaluate the similarity between the finalized DOME Copilot-generated annotations and the corresponding human manually curated annotations, a set of traditionally used metrics was employed to capture different aspects of textual similarity. This evaluation was conducted across the remaining subset of the benchmark dataset annotations ($n$ = 192) not utilized during the refinement process between prototype and finalized models. These metrics assess both lexical overlap and semantic similarity, allowing for a more comprehensive evaluation of the generated text. Specifically, Bilingual Evaluation Understudy (BLEU) (Papineni *et al.* 2002), Recall-Oriented Understudy for Gisting Evaluation (ROUGE-L) (Lin 2004), Metric for Evaluation of Translation with Explicit Ordering (METEOR) (Banerjee and Lavie 2005), and Bidirectional Encoder Representations from Transformers score (BERTScore) (Zhang *et al.* 2019) were used to provide complementary insights on the relationship between the DOME Copilot-generated and human ground-truth annotations. These metrics range from strict n-gram matching, which is typically more suited to LM-to-LLM evaluations, to more contextually aware semantic comparisons. BERTScore's semantic approach is most applicable for human-to-LLM evaluations, as the generative model is not attempting to reproduce identical phrasing as might be expected in stricter LLM output computational evaluations. A summary of these metrics, including their primary function and underlying computational approach, is presented in Table 2.1.

**Supplementary Table 2.1. Evaluation metrics summary.** Presentation of metrics alongside a description of their main function.

| Metric | Function | Method | Package used |
|---|---|---|---|
| BLEU | Rates the quality of the choices of words | Compares n-grams and counts the number of exact matches, applying a brevity penalty to discourage overly short outputs. | https://pypi.org/project/evaluate/ |
| ROUGE-L | Captures the structural coherence of the generated text | Compares the length of the longest common subsequence (LCS) of words, rewarding correct sequential order regardless of gaps. | https://pypi.org/project/evaluate/ |
| METEOR | Assesses text semantically and lexically | Aligns text using exact word matches, stemming, synonyms, and paraphrastic equivalence. | https://pypi.org/project/evaluate/ |
| BERTScore | Assesses text semantically | Computes pairwise cosine similarities using contextualized embeddings to match tokens to their most semantically similar counterpart. | https://github.com/Tiiiger/bert_score |

**BLEU**

The BLEU score was employed to assess the precision of the output by measuring how many n-grams (sequences of one or more words) in the generated text appear in the reference text. In principle, the

BLEU score rates the fluency of the output and the quality of the word choices. To discourage overly short outputs, BLEU incorporates a brevity penalty, which ensures that a generated sentence must not only match reference n-grams but also approximate the expected length. The BLEU score ranges from 0 to 1, achieving its maximum value of 1 only when all modified n-gram precisions are equal to 1 and no brevity penalty is applied, indicating that the generated text is an exact match to the reference

**ROUGE-L**

To evaluate the structural coherence of the generated text, ROUGE-L was implemented. This metric measures text similarity based on the length of the Longest Common Subsequence (LCS). The LCS is the longest series of words that appear in the same order in both the generated and reference texts, regardless of gaps. Unlike strict n-gram matching, the LCS does not require consecutive matches. Instead, it rewards sequences of words that appear in the same order in both texts, even if other words intervene. ROUGE-L assesses whether the output preserves the logical order of ideas or key phrases present in the reference, even when the exact wording differs. As with BLEU, ROUGE-L ranges from 0 to 1, reaching its maximum value of 1.0 when the LCS spans the full length of both the reference and generated text, meaning the two texts are identical in sequence.

**METEOR**

Semantic and lexical alignment was evaluated using METEOR. This is a recall-oriented and semantically sensitive metric that aligns the generated text with the reference using not only exact word matches but also matches based on stemming, synonyms, and paraphrastic equivalence. This approach makes METEOR more robust to wording variations and better correlated with human judgements. It also ranges from 0 to 1, attaining a maximum value of 1 when the generated text fully matches the reference in both content and ordering.

**BERTScore**

Finally, BERTScore was utilized to conduct a deeper assessment of the semantic similarity between the generated text and the reference. This metric leverages contextualized token embeddings from pre-trained transformer models. Instead of relying on exact n-gram overlap, BERTScore computes pairwise cosine similarities between tokens in the candidate and reference texts, greedily matching each token to its most semantically similar counterpart. BERTScore is highly robust to wording variations and better aligned for human text comparisons. It ranges from 0 to 1, reaching its maximum value when the generated text is semantically identical to the reference. Precision and recall are computed at the token level and are combined to calculate the F1 score.

To implement BERTScore in the evaluation pipeline, two different pretrained transformer backbones were tested to assess the trade-off between computational cost and evaluation accuracy. Specifically, the *microsoft/deberta-v3-base* (He, Gao, and Chen 2021) model and its larger *microsoft/deberta-xlarge-mnli* version were evaluated. The deberta-v3-base model provides a lighter configuration that enables faster inference while still producing high-quality contextual embeddings, whereas the deberta-xlarge-mnli model contains substantially more parameters, potentially allowing it to capture more subtle semantic relationships between texts.

Both models were initially used to compute the BERTScore for the benchmark prototype dataset to compare their behaviour in this evaluation setting. Processing the full set of entries with the deberta-v3-base model required approximately 3 hours, while the deberta-xlarge-mnli model required approximately 7.5 hours, reflecting its higher computational complexity. Although the larger model produced slightly higher similarity scores, the improvement was relatively small compared to the substantial increase in runtime. Consequently, the deberta-v3-base model was selected for the final analysis, providing an efficient balance between computational efficiency and reliable semantic similarity estimation. Based on this evaluation, the lighter model was used directly for all subsequent analyses of the finalized benchmark dataset.

**Metric Results**

The distributions of all evaluation metrics across the DOME annotation categories are shown in Supplementary Figure 2.1. Compared with BERTScore, the lexical and semi-semantic metrics (BLEU, ROUGE-L, and METEOR) exhibit lower absolute values and greater variability across entries. This behaviour is expected as these metrics rely more heavily on direct lexical overlap, and word order between the generated and reference annotations. They are therefore more sensitive to paraphrasing and stylistic variation in the DOME Copilot-generated text. Across the different DOME fields, the overall distributions remain broadly comparable. No single category displays systematically lower scores across all metrics. Among the three metrics, METEOR generally achieves higher values than BLEU and ROUGE-L. This reflects its ability to capture partial semantic matches through stemming and synonym alignment. Overall, these results provide complementary context to the BERTScore analysis. They illustrate the expected behaviour of overlap-based metrics when evaluating paraphrased annotations.

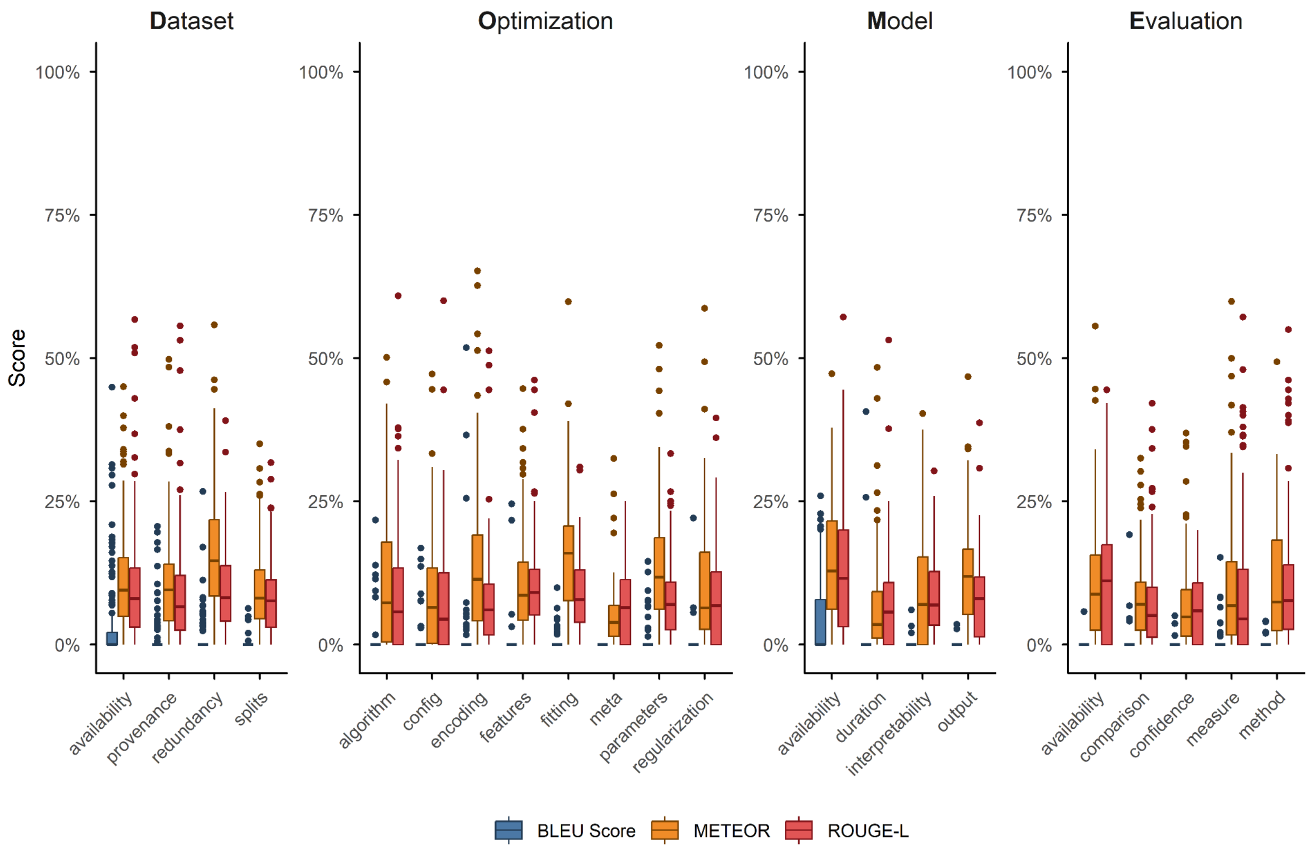


**Supplementary Figure 2.1. Distribution of the four evaluation metrics across DOME annotation fields.** Boxplots display the distribution of the BLEU, METEOR, and ROUGE-L metrics across the different methodology reporting fields. These scores compare the finalized DOME Copilot-generated annotations against the human ground truth from the benchmark dataset subset (*n* = 192). Each box represents the interquartile range of the scores across the registry entries. The central line indicates the median value. Points located beyond the whiskers represent outlier scores. A distinct colour identifies each specific evaluation metric.

The mean values of the evaluation metrics across all DOME Registry entries for the prototype benchmark were BLEU = 0.007 ± 0.023, ROUGE-L = 0.065 ± 0.068, METEOR = 0.137 ± 0.106, and the main text detailed BERTScore = 0.443 ± 0.092. For the final DOME Copilot benchmark, the corresponding values were BLEU = 0.009 ± 0.028, ROUGE-L = 0.092 ± 0.088, METEOR = 0.113 ± 0.101, and BERTScore = 0.429 ± 0.080. As expected, the lexical overlap metrics generally yield lower scores compared with the embedding-based semantic metric. In particular, BLEU exhibits the lowest values among the evaluated metrics. This reflects its strict reliance on exact n-gram overlap between the generated and reference texts. DOME Copilot-generated annotations frequently paraphrase the reference descriptions rather than reproducing identical wording. Consequently, strict overlap metrics such as BLEU tend to underestimate similarity in this context. In contrast, METEOR and BERTScore incorporate mechanisms to capture semantic relationships between words. These metrics produce higher similarity scores and provide a more informative assessment of the semantic correspondence between the generated outputs and the human-curated annotations.

**AlphaFold 2: Gold-Standard Metric Validation Case**

To mitigate the potential limitations associated with large-scale automated comparisons against highly heterogeneous human-curated datasets, a gold-standard annotated manuscript was additionally evaluated. This validation case focused on the AlphaFold 2 publication (Jumper *et al.* 2021). This specific manuscript was manually curated by a domain expert curating for the DOME Registry. This expert curation ensures the ground-truth annotations represent a high-quality reference example with rigorously validated entries. The AlphaFold 2 annotations were evaluated across all reporting fields. The generative metrics achieved notably higher mean scores of BLEU = 0.016, ROUGE-L = 0.131, METEOR = 0.149, and BERTScore = 0.503. The manuscript represents a highly complex AI methodology spanning multiple modular components. The strong semantic similarity demonstrated by the BERTScore highlights two critical aspects of the finalized DOME Copilot performance. It first supports the relevant extraction and summarization capabilities of the model when processing highly complex architectural descriptions. Furthermore, it provides strong confidence in the automated evaluation methodology when comparing the AI generated outputs against a high-quality, expertly curated human baseline.

**References (Supplementary Material 2)**

Banerjee S, Lavie A. METEOR: an automatic metric for MT evaluation with improved correlation with human judgments. In: Goldstein J, Lavie A, Lin CY et al. (eds). Proceedings of the ACL Workshop on Intrinsic and Extrinsic Evaluation Measures for Machine Translation and/or Summarization. *Association for Computational Linguistics*; 2005, 65–72. https://aclanthology.org/W05-0909/

He P, Gao J, Chen W et al. DeBERTaV3: improving DeBERTa using ELECTRA-style pre-training with gradient-disentangled embedding sharing. *arXiv* [Preprint] 2021. https://doi.org/10.48550/arXiv.2111.09543

Jumper J, Evans R, Pritzel A et al. Highly accurate protein structure prediction with AlphaFold. *Nature* 2021;596:583–9. https://doi.org/10.1038/s41586-021-03819-2

Lin CY. ROUGE: a package for automatic evaluation of summaries. In: Text Summarization Branches Out. *Association for Computational Linguistics*; 2004, 74–81. https://aclanthology.org/W04-1013/

Papineni K, Roukos S, Ward T et al. BLEU: a method for automatic evaluation of machine translation. In: Proceedings of the 40th Annual Meeting of the Association for Computational Linguistics. *Association for Computational Linguistics*; 2002, 311–8. https://doi.org/10.3115/1073083.1073135

Zhang T, Kishore V, Wu F et al. BERTScore: evaluating text generation with BERT. *arXiv* [Preprint] 2019. https://doi.org/10.48550/arXiv.1904.09675
